# Htm1p-Pdi1p is a folding sensitive mannosidase that marks N-glycoproteins for ER-associated protein degradation

**DOI:** 10.1101/049825

**Authors:** Yi-Chang Liu, Danica Galonić Fujimori, Jonathan S. Weissman

## Abstract

Our understanding of how the endoplasmic reticulum-associated protein degradation (ERAD) machinery efficiently targets terminally misfolded proteins while avoiding the misidentification of nascent polypeptides and correctly folded proteins is limited. For luminal N-glycoproteins, demannosylation of their N-glycan to expose a terminal α1,6-linked mannose is necessary for their degradation via ERAD, but whether this modification is specific to misfolded proteins is unknown. Here we report that the Htm1p-Pdi1p complex acts as a folding-sensitive mannosidase for catalyzing this first committed step. We reconstitute this step *in vitro* with Htm1p-Pdi1p and model glycoprotein substrates whose structural states we can manipulate. We find that Htm1p-Pdi1p is a glycoprotein-specific mannosidase, which preferentially targets nonnative glycoproteins trapped in partially structured states. As such, Htm1p-Pdi1p is suited to act as a licensing factor that monitors folding in the ER lumen and preferentially commits glycoproteins trapped in partially structured states for degradation.

## Introduction

Proteins destined for the secretory pathway enter the endoplasmic reticulum (ER) in unfolded states and generally leave only after they have reached their native conformation. Protein folding in the ER is challenging and often error-prone [1, 2]. Thus the ER must continuously monitor the pool of folding proteins and remove terminally misfolded forms before they induce toxicity. This is accomplished through a series of pathways collectively referred to as the ER-associated protein degradation (ERAD) machinery [3]. Generally, the process of ERAD involves three steps: 1) identification of the misfolded protein, 2) retrotranslocation and polyubiquitination of the misfolded protein through individual E3 ubiquitin ligase complexes, and 3) degradation of the misfolded protein by the ubiquitin-proteasome system in the cytosol.

Each ERAD pathway needs to accurately identify substrates to avoid overly promiscuous destruction of functional proteins while preventing the accumulation of toxic forms [1]. For the ERAD pathway responsible for the destruction of proteins with misfolded luminal domains (ERAD-L) [4, 5], this process of discrimination depends not only on the folding status of a polypeptide but also on its N-glycosylation state [6-11]. Upon entering the ER lumen, nascent proteins acquire en bloc a Glc_3_Man_9_GlcNAc_2_ high-mannose oligosaccharide, appended to a specific subset of asparagine residues (Fig. 1*A*). The N-glycan is then deconstructed monosaccharide-by-monosaccharide to Man_8_GlcNAc_2_ (Man8) in the ER by a series of glycosidases concurrently with the protein folding process [12, 13]. The action of the mannosidase Mns1p, which is responsible for generating Man8, has been proposed to yield a “timer” that universally gives all N-glycoproteins a defined period of time to fold during which they are immune from ERAD-L [8, 12]. Misfolded proteins that are destined for degradation via ERAD-L are further demannosylated by Htm1p (also known as Mnl1p), an enzyme that removes the α1,2-linked mannose from the C branch of Man8, generating a Man_7_GlcNAc_2_ (Man7) glycan structure with a terminally exposed α1,6-linked mannose that serves as a signal for ERAD-L commitment [14, 15]. The Yos9p lectin subsequently binds this terminally exposed α1,6-linked mannose and, in coordination with Hrd3p, queries the misfolded regions on potential substrates [6, 7, 9-11, 16]. Only the presence of both the Htm1p-generated N-glycan signal and the misfolded regions on the substrate enables Yos9p-Hrd3p to trigger the downstream retrotranslocation, polyubiquitination, and proteasomal degradation of target proteins [7, 14, 17]. The physiological importance of these checkpoints is exemplified by the severe growth defect acquired when the requirement for Hrd3p and Yos9p was bypassed by overexpression of *HRD1*, which encodes the downstream E3 ubiquitin ligase for this pathway [6].

**Figure 1.**
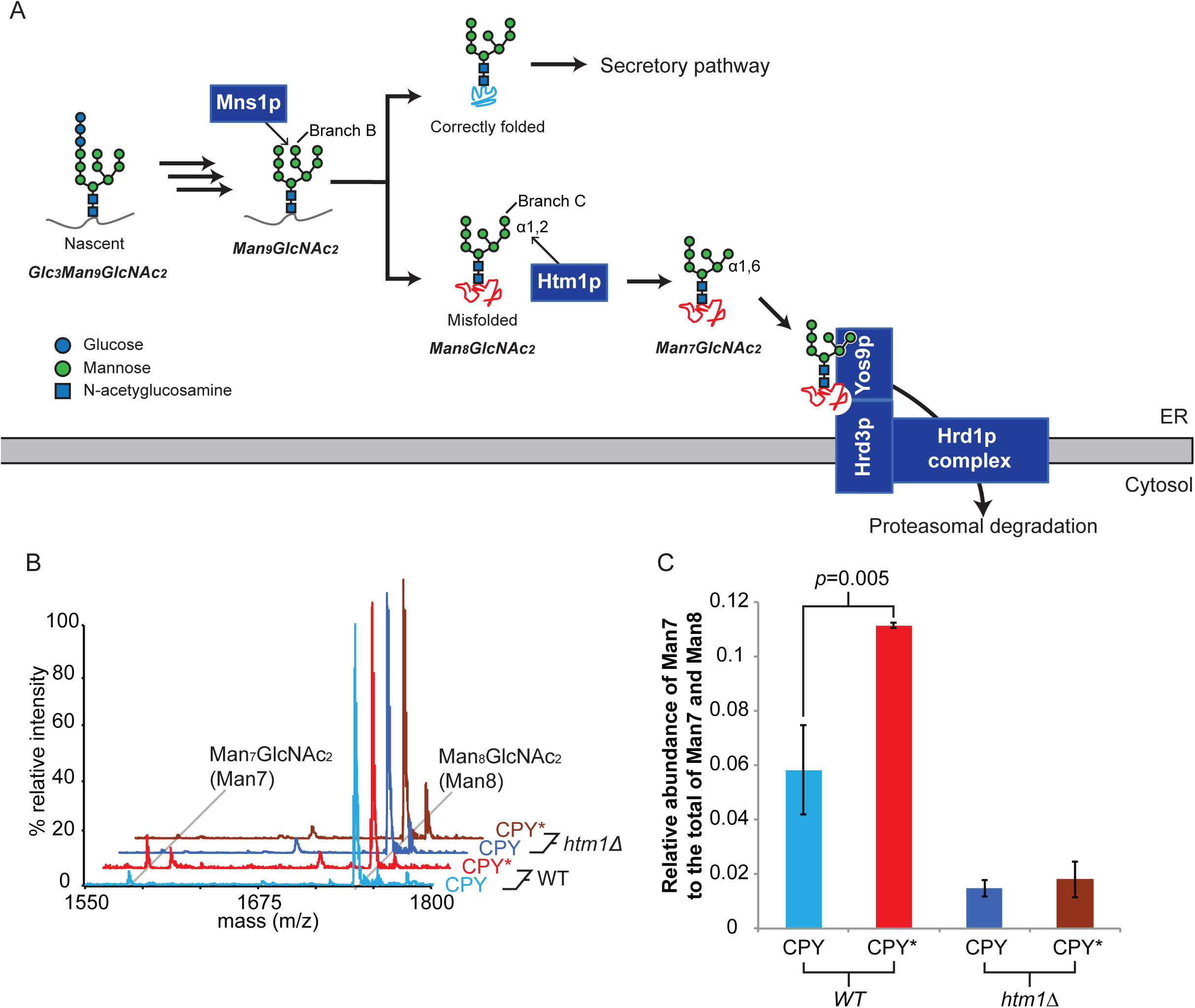
HTM1 mediates N-glycan processing for ERAD-L commitment. **(*A*)** Scheme of N-glycan processing and ERAD-L commitment. Upon entering the ER, nascent N-glycoproteins acquire the Glc_3_Man_9_GlcNAc_2_ *en bloc*, which is step-by-step deglycosylated as the nascent polypeptide folds. Mns1p catalyzes the last universal deglycosylation step to generate the Man_8_GlcNAc_2_ glycan. While native proteins are targeted for downstream secretory pathway, misfolded proteins are targeted by Htm1p, which removes the α1,2-linked mannose at the C branch from the Man_8_GlcNAc_2_ glycan. The resulting terminally exposed α1,6-linked mannose and misfolded regions on the polypeptide chain are then read by Yos9p. Along with Hrd3p, Yos9p then permits the downstream retrotranslocation and polyubiquitination through the Hrd1p transmembrane complex for eventual proteasomal degradation. **(*B*)** Examples of glycan profiles of CPY and CPY* expressed in wild type (*WT*) or *htm1*Δ yeast strains as measured by MALDI-TOF MS after the glycans were released from CPY/CPY* by PNGase F treatment. Each glycan was assigned to the m/z corresponding to its predicted [M+Na]^+^ value. **(*C*)** The relative abundance of Man7 is higher in CPY* than CPY in wild-type yeast, which is dependent on *HTM1*. The relative MS peak intensity of Man7, as found in (b), is normalized to by the total of Man7 and Man8 as a proxy for its relative abundance. Data shown are mean values ± one standard deviation (s.d.) from biological triplicates. The *p* value was calculated by unpaired *t* test.

As the first committed step, the generation of the terminally exposed α1,6-linked mannose by Htm1p has the potential to determine which proteins, and at what stage during their folding process, are shunted down the ERAD-L pathway. This potential “licensing” role has been supported by *in vivo* observations that genetic deletion of *HTM1*, as well as point mutations in its putative active sites, not only retards the degradation of misfolded N-glycoproteins, but also decreases substrate-binding by Yos9p [11, 14, 17-19]. This retardation of ERAD by deletion of *HTM1* can be circumvented in a genetic background that generates the terminally exposed α1,6-linked mannose [7]. The importance of the mannosidase function of Htm1p is further evidenced by the conserved role of HTM1 homologs for the N-glycan-dependent ERAD pathway [20]. Finally, both *in vivo* and *in vitro* studies have provided evidence for the mannosidase activity of Htm1p in generating Man7 [14, 15, 21]. However, despite the well-documented role of Htm1p in the N-glycan processing for ERAD-L commitment, it is still unclear how specifically and accurately Htm1p targets misfolded proteins. This uncertainty is mainly due to the challenges in simultaneously monitoring protein conformations and their N-glycosylation states in the highly heterogeneous pool of glycoproteins in the ER, as well as the presence of multiple competing ER quality control pathways [22]. Additionally, only minimal mannosidase activities of Htm1p were observed in previous *in vitro* studies [15, 21]. To advance our understanding of this commitment step, it is critical to develop a reconstituted system for direct investigation of enzymatic activities of Htm1p against glycoproteins with defined glycosylation and folding states [23].

Here, we reconstituted this ERAD-L commitment step, which consists of a recombinantly expressed Htm1p-Pdi1p complex and glycoprotein substrates with various native and nonnative conformations. Our results reveal that Htm1p-Pdi1p is a glycoprotein-specific mannosidase complex that preferentially demannosylates intrinsically or artificially misfolded proteins compared to their native counterparts. Furthermore, among the various nonnative conformations, Htm1p-Pdi1p prefers partially structured proteins over globally unfolded ones. Our findings suggest that Htm1p-Pdi1p monitors the protein folding states in the ER lumen and preferentially licenses glycoproteins trapped in partially structured states for ERAD-L.

## Results

### Generation of Man7 *in vivo* is folding-and *HTM1*-dependent

It has been reported that the prototypic ERAD-L substrate, CPY*, contains higher levels of Man7 than its folding-competent counterpart, pro-carboxypeptidase Y (CPY) in a wild-type yeast background [11]. We first explored whether the higher level of Man7 of CPY* reflects the differences in intrinsic folding competence between CPY* and CPY, or rather results passively from the prolonged ER retention of CPY*. Utilizing a previously described overexpression construct [24], we expressed and purified native CPY and CPY* that are retained in the ER by a C-terminal HDEL tag. We used MALDI-TOF mass spectrometry to determine the glycan profile of CPY and CPY*. This approach provides higher throughput capacity than HPLC-based measurement while still allowing quantitative analysis of neutral glycans [25]. As such, it is also amenable to the analysis of mannosidase activities described below. Both CPY and CPY* purified from wild-type (BY4741) yeast strain carry predominantly Man8 and minor populations of Man7, Man9 and Hex_10_GlcNAc_2_ (Hex10), which is likely GlcMan_9_GlcNAc_2_ (Fig. S1 *A* and *B*). When CPY and CPY* were purified from a *mns1*Δ*htm1*Δ background, the dominant glycan species shifted from Man8 to Man9, verifying that the majority of Man8 from the WT strain is the product of Mns1p, the putative substrate of Htm1p (Fig. S1 *A* and *B*). Importantly, CPY* carries a significantly higher level of Man7 than CPY (Fig. 1 *B* and *C*). The majority of the signal of Man7 disappeared when CPY and CPY* were purified from the *htm1*Δ strain background. This observation suggests that despite both being retained in the ER, CPY* and CPY are differentially demannosylated to Man7 in an *HTM1*-dependent manner. Given that CPY and CPY* are only different at the G255R point mutation on CPY* that disrupts the folding of the hydrophobic core of native CPY [26], we conclude that the generation of Man7 in the ER is correlated with the intrinsic folding competence of the underlying protein.

### Reconstitution of an Htm1p-Pdi1p complex

In order to explore the activity of Htm1p *in vitro*, we tested the feasibility of producing Htm1p from the native host *Saccharomyces cerevisiae*. We introduced a 3xFLAG tag, followed by a HDEL tail before the stop codon, to the C-terminus of Htm1p for affinity purification. The chromosomally tagged version is similarly functional at degrading CPY* as the wild-type Htm1p (Fig. 2*A*). While the endogenous level of Htm1p is low, we can enhance the expression of Htm1p by chromosomally substituting its endogenous promoter with a *TDH3* promoter (Fig. 2*B*). Consistent with a previous study [27], Htm1p readily forms a disulfide complex with endogenous Pdi1p as resolved by non-reducing SDS-PAGE (Fig. 2*B*). Large-scale immunoprecipitation yielded an Htm1p-Pdi1p complex with near 1:1 stoichiometry (Fig. 2*C*), with a yield of 200 μg of the complex from three liters of yeast culture. Endoglycosidase H treatment of the complex confirmed that both Htm1p and Pdi1p are N-glycosylated, indicating that they are correctly targeted to the secretory pathway (Fig. 2*D*). The complex behaved as a monodisperse species in size-exclusion column chromatography (Fig. 2*E*), which contained a mixture of both disulfide-linked and non-covalent complexes of Htm1p and Pdi1p (Fig. 2*ƒ*). We did not detect any monomeric Htm1p or Pdi1p corresponding to their predicted molecular weight (90.2 kDa for Htm1p, and 56.0 kDa for Pdi1p). Finally, we observed the mannosidase activity against the most preferred glycoprotein substrate, RBsp, only in the fractions containing the Htm1-Pdi1p complex (Fig. 2*G*), *vide infra*.

**Figure 2.**
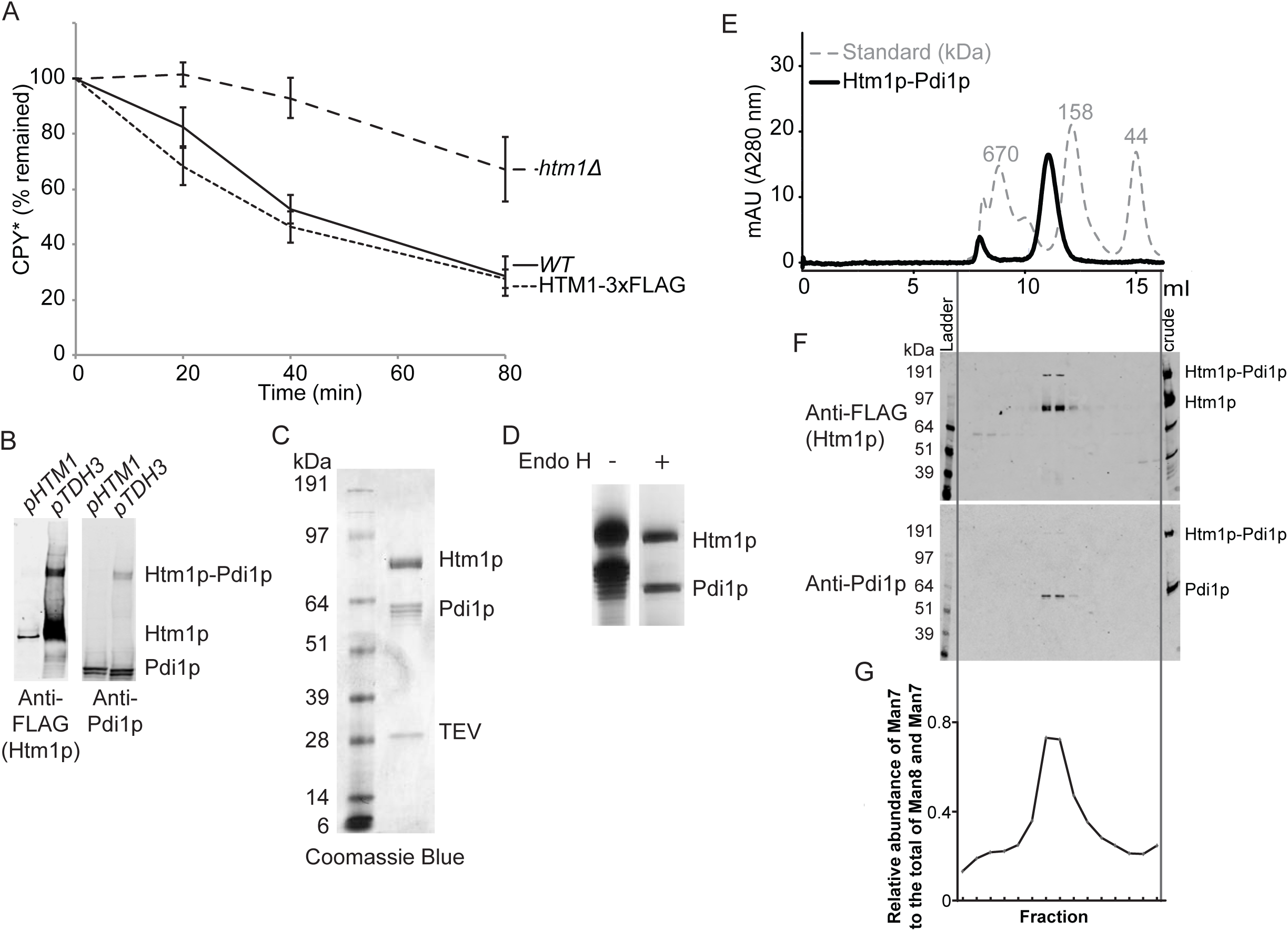
Recombinant preparation of Htm1-Pdi1p from *S. cerevisiae*. **(*A*)** Degradation of HA-tagged CPY* is the same in a wild-type BY4741 (WT) and chromosomally 3xFLAG-HDEL-tagged *HTM1* (HTM1-tag). Equal amount of cells growing at mid-log phase were harvested, subjected to reducing SDS-PAGE, and quantitative western blot for the HA signal at each time point relative toe zero time point. Data shown are the mean values ± one standard error of the mean measured from four biological replicates. **(*B*)** Expression levels of Htm1p-3xFLAG driven by either endogenous *HTM1* promoter (*pHTM1*) or chromosomally inserted *TDH3* promoter (*pTDH3*) as checked by non-reducing SDS-PAGE and western blot using anti-FLAG (mouse) and anti-Pd1ip (rabbit) primary antibodies and corresponding secondary IR-fluorescent antibodies. **(*C*)** Specific interaction between Htm1p and Pdi1p after large-scale immunoprecipitation with anti-FLAG affinity resin followed by TEV protease treatment to release Htm1p. Shown is a Coomassie Blue R250-stained gel image after reducing SDS-PAGE. **(*D*)** The N-glycosylation states of Htm1p and Pdi1p as checked by endoglycosidase H (Endo H) treatment. **(*E*)** Superdex 200 10/300GL size exclusion column chromatography of Htm1p-Pdi1p eluted from anti-FLAG resin by 3xFLAG peptide. **(*ƒ*)** Western blot analysis for the distribution of Htm1p-3xFLAG and Pdi1p after non-reducing SDS-PAGE of the fractions separated by the size-exclusion column chromatography. **(*G*)** The relative abundance of Man7 in the total of Man7 and Man8 on RBsp after one-hour incubation with individual fractions collected from the size-exclusion column chromatography of purified Htm1p-Pdi1p.

### Htm1p-Pdi1p preferentially demannosylates CPY* and misfolded CPY variants

We first explored the mannosidase activity of reconstituted Htm1p-Pdi1p against CPY and CPY* that we purified from wild-type yeast. After a 20-hour reaction, Htm1p-Pdi1p converted a portion of Man8 on CPY and CPY* to Man7 (Fig. 3 *A* and *B*). Consistent with our *in vivo* observation, the increase in Man7 on CPY* is significantly higher than that on CPY after incubation with Htm1p-Pdi1p (Fig. 3*C*). To further validate that Htm1p-Pdi1p preferentially targets nonnative proteins, we reductively denatured CPY to remove its structurally essential disulfide bonds [28]. Reduced CPY was then oxidized under denaturing condition into “scrambled” species with randomly distributed nonnative disulfide bonds (Scr-CPY), or was carbamidomethylated to block reformation of disulfide bonds (Carb-CPY). Htm1p-Pdi1p also demannosylated both Scr-CPY and Carb-CPY more efficiently than the native form (Fig. 3*D*). Taken together, these results suggest that our reconstituted Htm1p-Pdi1p is an active mannosidase and preferentially acts on intrinsically folding-incompetent CPY* and artificially misfolded CPY variants compared to their native counterpart.

**Figure 3.**
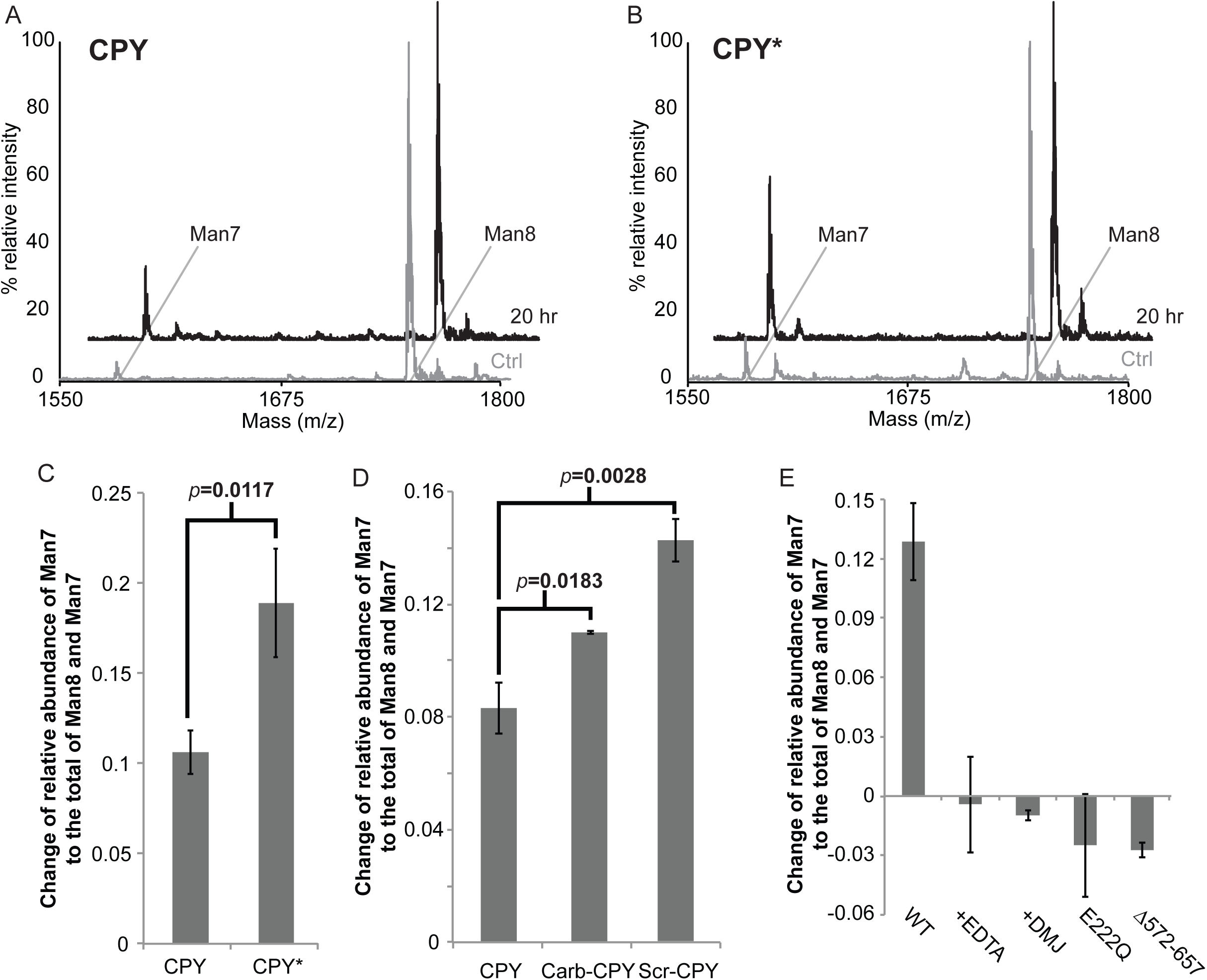
Htm1p-Pdi1p preferentially demannosylates CPY* and nonnative CPY variants. (***A* and *B***) Examples of glycan profiles of CPY **(a)** and CPY* **(b)** before and after twenty-hour incubation with Htm1p-Pdi1p. For the experiment, 2 μM CPY or CPY* was incubated with 0.1 μM Htm1p-Pdi1p at 30°C for twenty hours. The reaction was stopped with 25 mM EDTA. CPY or CPY* was then separated from Htm1-Pdi1p by SDS-PAGE for subsequent in-gel glycan release and MALDI-TOF MS analysis. **(*C*)** Htm1p-Pdi1p generates more Man7 on CPY* than CPY after reaction. The mannosidase activity is measured based on the change of relative abundance of Man7 on CPY and CPY* after twenty-hour incubation with Htm1p-Pdi1p. Data shown are the mean values ± one s.d. from experimental triplicates. The *p* value was calculated by unpaired *t* test. **(*D*)** Htm1p-Pdi1p generated more Man7 on Carb-CPY and Scr-CPY than CPY. Numbers shown are the mean ± one s.d. from experimental triplicates. The *p* value was calculated by unpaired *t* test. (***E***) The mannosidase activity of Htm1p-Pdi1p against CPY* was inhibited by EDTA (+EDTA), 1-deoxymannojirimicin (+DMJ), mutations including E222Q and Δ572-657. Quantification was carried out as in (c). Data shown are the mean values ± one s.d. from experimental triplicates.

We then wanted to verify whether demannosylation is catalyzed through the predicted mannosidase domain of Htm1p, and whether Pdi1p is required for this process. We carried out the reaction in the presence of EDTA, which sequesters the structurally essential Ca^2+^, as well as 1-deoxymannojirimycin (DMJ), which inhibits the GH47 mannosidase family that Htm1p is predicted to belong to [29]. Both chemicals inhibited the generation of Man7 on CPY*(Fig. 3*E*). We further verified the predicted mannosidase function of Htm1p with an Htm1 mutant with one of the putative active site residues Glu_222_ mutated to Gln (E222Q) [14, 30], which was indeed incapable of generating Man7 on CPY* (Fig. 3*E*). Finally, we made a second Htm1 mutant that has a truncation of the amino acyl region from 572 to 657 (Δ572-657), which covers the region necessary for interaction with Pdi1p (Fig S2*A*) [14, 21, 27]. Immunoprecipitation confirmed that Δ572-657 showed little detectable interaction with Pdi1p (Fig S2*B*). Similar to the catalytically dead E222Q, Δ572-657 was incapable of generating Man7 on CPY* (Fig. 3*E*). Collectively, our results support that the predicted mannosidase domain of Htm1p mediates the demannosylation reaction, and that the interaction with Pdi1p is required for this activity.

The conversion of Man8 to Man7 on CPY* continued during prolonged incubation with Htm1p-Pdi1p (Fig. S3*A*), but this rate of demannosylation is nonetheless much slower than what would be expected from the *in vivo* half-life of CPY*, which is about 60 minutes (Fig. 2*A*). This slow rate in principle could be a consequence of the fact that CPY has four N-glycans, and thus Htm1p-Pdi1p may only act on a subset of glycans on CPY*. Indeed, the C-terminal most N-glycan of CPY* is known to be necessary and sufficient to support its ERAD *in vivo* [17, 31, 32]. However, we found that *in vitro*, Htm1p-Pdi1p demannosylated a CPY* mutant, in which this ERAD-competent glycosylation site, Asn_479_, was mutated to Gln (CPY*1110), to a level similar to CPY* with all four glycosylation sites (Fig. S3*B*). A similar level of demannosylation was observed on another CPY* mutant in which the other three glycosylation sites (Asn_124_, Asn_198_, and Asn_279_) were all mutated to Gln (CPY*0001). The glycosylation site is thus not likely the key factor limiting the catalytic activity. Increasing the amount of Htm1p-Pdi1p by four folds did not proportionally increase the yield, suggesting that the enzyme concentration is not the limiting factor (Fig. S3*E*).

The incomplete reaction may also reflect conformational heterogeneity or the aggregation-prone nature of CPY*. In our hand, the majority of purified CPY* is in oligomeric forms (Fig. S4), and we observed formation of visible aggregates from both CPY* and artificially misfolded CPY variants after incubation with Htm1p-Pdi1p for twenty hours. A potential consequence of aggregation is that the glycans become inaccessible to the deep active pocket of Htm1p as predicted by the GH47 family mannosidase domain [33]. First, we explored whether Htm1p-Pdi1p has higher activity against free Man8 glycan. Htm1p-Pdi1p only generated limited amount of Man7 after 24 hours of reaction (Fig. S5*A*). In contrast, Mns1p, which similarly contains to a GH47 family mannosidase domain, was able to completely demannosylate Man9 into Man8 (Fig. S5*B*). To further assess the accessibility to glycans on CPY, we tested the activity of Mns1p against Man9-carrying CPY (CPYm9) and its scrambled form (Scr-CPYm9) purified from the *mns1*Δ*htm1*Δ strain background. Mns1p converted most of Man9 on both CPYm9 and Scr-CPYm9 into Man8, suggesting that the glycans are accessible on both native and nonnative forms of CPY (Fig. 4). Mns1p alone did not produce any detectable Man7. Only co-treatment of Mns1p and Htm1p-Pdi1p yielded Man7 to a level similar to what we observed in Figure 3. We found that Htm1p-Pdi1p alone, without Mns1p, was also capable of demannosylating Man9 on CPYm9 and Scr-CPYm9 into Man8, albeit to a lower extent. Similar to the results in Figure 3*D*, the level of Man8 generated by Htm1p-Pdi1p treatment was higher on Scr-CPYm9 than CPYm9. Collectively, these observations suggest intrinsic differences in substrate specificity between Mns1p and Htm1p-Pdi1p: while Mns1p is capable of targeting both free glycans and glycoproteins independently of the attached protein conformations, Htm1p-Pdi1p preferentially targets glycans installed on nonnative proteins. In addition, our findings suggest that Htm1p-Pdi1p is capable of bypassing the action of Mns1p to directly demannosylate Man9, which is further supported by previous *in vivo* studies [10, 30, 34].

**Figure 4.**
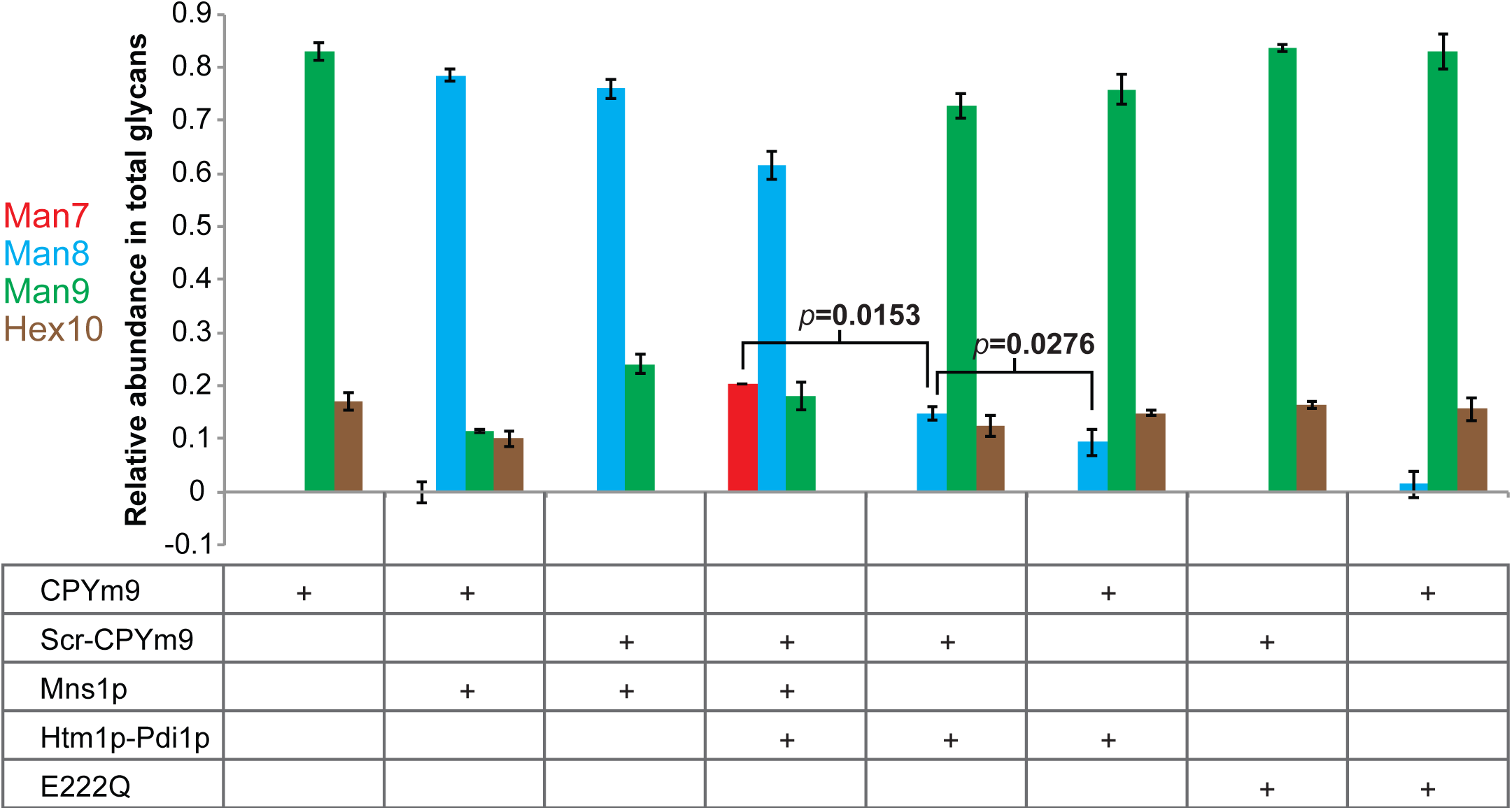
Htm1p-Pdi1p demannosylates α1,2-Linked Mannose from Branch C of Man9. The relative abundance of each glycan on CPYm9 or Scr-CPYm9 after twenty-hour incubation with Mns1p and/or Htm1p-Pdi1p, or E222Q. Data shown are the mean values ± one s.d. from experimental triplicates. The *p* value was calculated by unpaired *t* test.

### Htm1p-Pdi1p preferentially demannosylates partially structured RNase B variants

To this point, our findings support two potential models that can describe how Htm1p-Pdi1p generates the α1,6-lniked mannose signal for ERAD-L commitment. In one model, the catalysis by Htm1p-Pdi1p may be intrinsically slow and stochastic as previously suggested [15, 21], and unfolding facilitates demannosylation by increasing access to the glycan. Alternatively, Htm1p-Pdi1p may preferentially target specific folding states of nonnative proteins. To test these two potential models, we introduced bovine pancreatic ribonuclease B (RNase B) as an alternative substrate. RNase B is identical to the well-characterized classic protein folding substrate RNase A protein with the exception that it contains a single N-glycosylation site [35]. The folding of RNase B has been extensively studied, and the proteins can be manipulated to form well-defined, homogeneous states of native, misfolded or unfolded conformations [36-38]. Using a concanavalin A lectin-based enrichment approach [39], we can enrich for Man8-abundant RNase B from commercial sources in biochemical quantities (Fig. S6). To generate a nonnative variant of RNase B, we proteolytically removed the N-terminal S peptide from native RNase B to produce RNase BS protein (RBsp) (Fig. 5*A*). RBsp is trapped in a compact, disordered but nonetheless well-behaved state, and it can be readily reconstituted into a native-like RNase BS form by addition of the S peptide [37].

**Figure 5.**
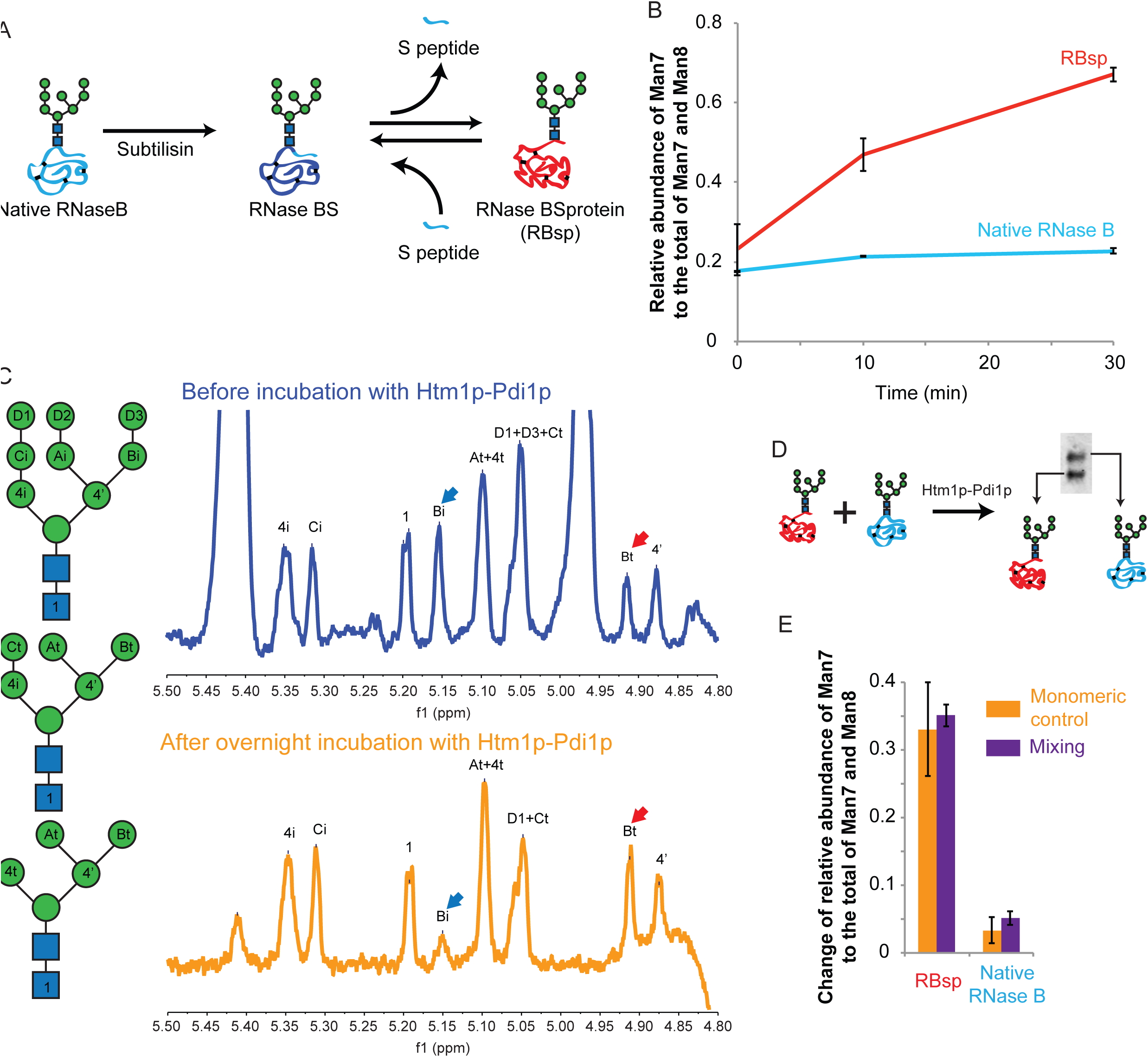
Figure 5. Htm1p-Pdi1p efficiently demannosylates RBsp. **(*A*)** Schematic presentation of the generation of RBsp. Native RNase B was treated with subtilisin in a 100:1 ratio at 4°C for twenty-four hours to generate RNase BS with a nick between the N-terminal S peptide and the rest of the protein (RBsp). RNase BS was then subjected to precipitation with 10% Trichloroacetic acid (TCA) to remove the soluble S peptide from the precipitated RBsp. **(*B*)** Change of the relative abundance of Man7 on RBsp and native RNase B during a time course incubation with Htm1p-Pdi1p. Data shown are the mean values ± one s.d. from experimental triplicates **(*C*)** ^1^H-NMR spectra (D_2_O, 400 MHz) of the glycan released from RNase B before and after overnight incubation with Htm1p-Pdi1p. Note the change of the height of the peaks corresponding to the α1,6-linked mannose on the Branch C with (Bi, blue arrow) and without (Bt, red arrow) the terminal α1,2-linked mannose (D3) after incubation with Htm1p-Pdi1p. Shown are the regions covering the H-1 on the anomeric carbon of each monosaccharide after solvent suppression by the presaturation method. The peak assignment were based on published data [40], using symbols shown in the left scheme. **(*D*)** Schematic presentation of the co-reaction of RBsp and native RNase B with Htm1p-Pdi1p. RBsp and native RNase B were pre-mixed at 1:1 ratio and incubated with Htm1p-Pdi1p for twenty minutes, and subsequently separated by SDS-PAGE for MS analysis of the individual glycan profile. **(*E*)** Change of the relative abundance of Man7 on premixed RBsp and native RNase B after incubation with Htm1p-Pdi1p for twenty minutes. Data shown are the mean values ± one s.d. from experimental triplicates

Remarkably, Htm1p-Pdi1p differentiated between native RNase B and RBsp, and only efficiently demannosylated RBsp (Fig. 5*B*). Demannosylation of Man8 on RBsp into Man7 nearly reached the plateau after one hour (Fig. S7*A*). Consistent with the results in Figure 4, we observed the disappearance of Man9 on RBsp during the reaction with Htm1p-Pdi1p (Fig. S7 *A* and *B*). The residual Man8 can be attributed to the other two less abundant Man8 isoforms on RNase B that already have the C branch α1,2-mannose removed [40]. Indeed, all of the Man9 and Man8 on RBsp were removed after co-treatment with Mns1p and Htm1p-Pdi1p (Fig. S7*B*), which also verified that Mns1p and Htm1p-Pdi1p target different mannoses on Man9. We further explored whether Htm1p-Pdi1p targets the α1,2-linked mannose on the Branch C or the Branch A, a question that has never been thoroughly investigated. To obtain a structural view of which mannose is removed by Htm1p-Pdi1p, we performed ^1^H-NMR analysis on the glycans released from denatured RNase B before and after an overnight reaction with Htm1p-Pdi1p. Indeed, we found only the α1,2-linked mannose on the Branch C was removed after an overnight incubation with Htm1p-Pdi1p (Fig 5*C*).

We further investigated whether Htm1p-Pdi1p can be *trans*-activated by RBsp, or *trans*-inhibited by CPY*. We tested this possibility by incubating Htm1p-Pdi1p with CPY* for two hours, and then adding RBsp. Molecular weight differences between CPY* (76.7 kDa) and RBsp (13.2 kDa) allow them to be readily separated by SDS-PAGE for subsequent glycan profiling. No obvious differences was observed on demannosylation of RBsp in the presence of absence of CPY*, and *vice versa* (Fig. S8 *A* and *B*). We further extended this experiment with a mixture of native RNase B and RBsp, which can be similarly separated by SDS-PAGE given their difference in molecular weight by the presence or absence of the S-peptide (Fig. 5*D*). The presence of RBsp did not induce the demannosylation of native RNase B by Htm1p-Pdi1p (Fig. 5*E*). Collectively, these data establish that Htm1p-Pdi1p is not inhibited or activated by CPY* or RBsp *in trans*. Additionally, Htm1p-Pdi1p can distinguish its preferred substrates among a mixture of different glycoproteins.

To further characterize how accurately Htm1p-Pdi1p differentiates proteins with different nonnative conformations, we generated three additional variants of RNase B: 1) subtilisin-nicked non-covalent complex of RBsp and S-peptide (RNase BS), 2) disulfide-scrambled RNase B (Scr-RB), and 3) cysteine-carbamidomethylated RNase B (Carb-RB) (Fig. 6*A*). Consistent with the known properties of these variants [36, 37], our far-UV circular dichroism analysis verified that these RNase B variants cover a range of different conformations (Fig. 6*B*): RNase BS behaves essentially identically to native RNase B; RBsp is a compact folding intermediate with relatively abundant secondary structure features; Scr-RB consists of random, less compact structures with residual signals from secondary structures; Carb-RB consists of globally unfolded species with the majority of the signals from random coils. We further confirmed this increasing trend of unfolding from native RNase B to Carb-RB by monitoring the differences in peptide backbone flexibility and glycan accessibility of different RNase B variants through trypsin and PNGase F treatment (Fig. S9 *A* and *B*). Of a particular note, unlike CPY* and nonnative CPY variants, both Scr-RB and Carb-RB did not oligomerize or aggregate even after overnight incubation with Htm1p-Pdi1p at 30°C (Fig. S9 *C* and *D*).

**Figure 6.**
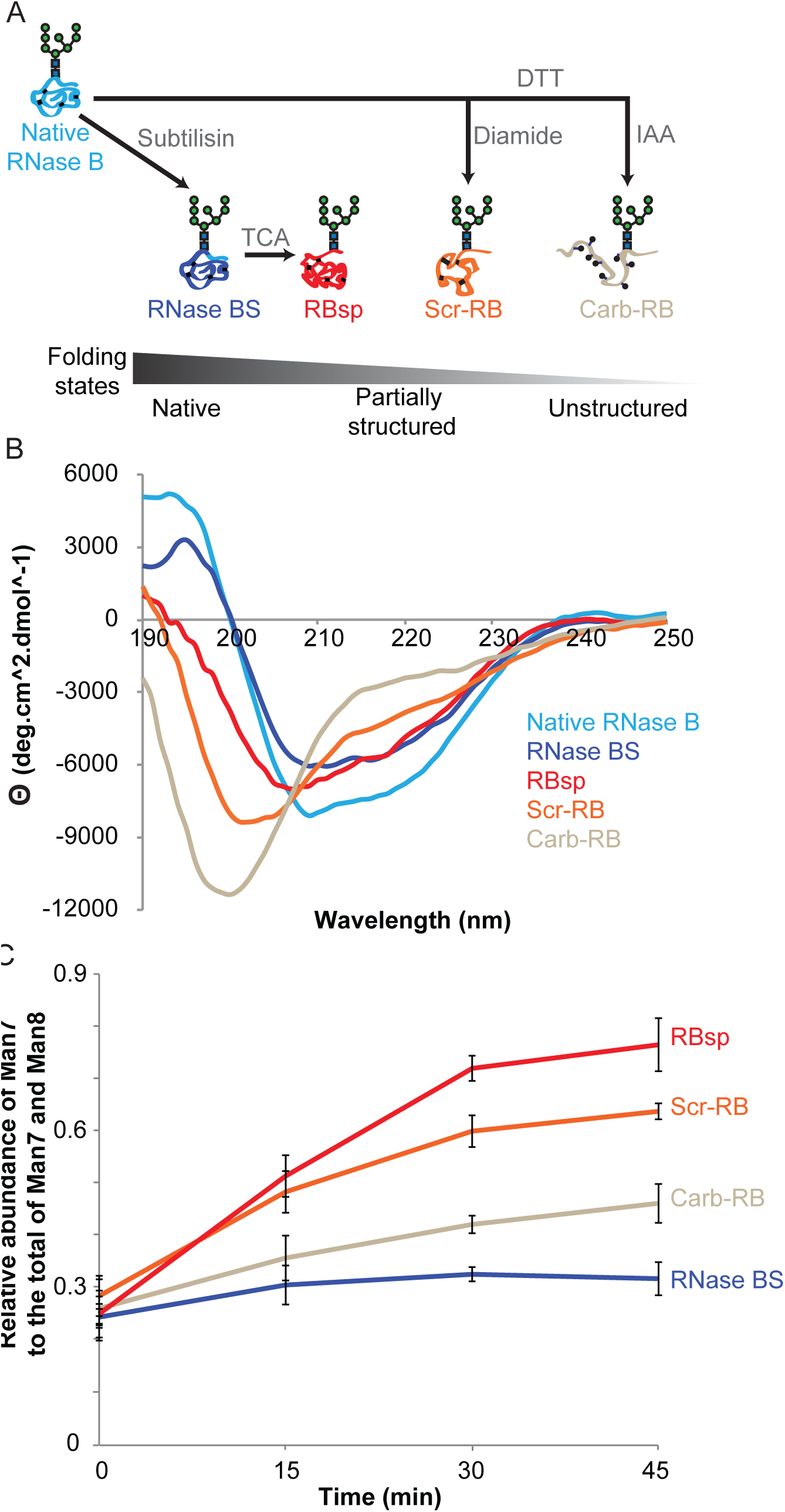
Htm1-Pdi1p preferentially demannosylates partially structured RNase B variants. **(*A*)** Schematic presentation of the generation of RNase B variants. RNase BS and RBsp are generated through subtilisin and TCA treatment as described in Fig.5a. Additionally, RNase B is reductively denatured by DTT in 6 M guanidine hydrochloride and then either disulfide-scrambled by diamide to produce Scr-RB or cysteine-carbamidomethylated by iodoacetamide (IAA) to produce Carb-RB. ***(B)*** Circular dichroism analysis of RNase B variants. ***(C)*** The relative abundance of Man7 on RNase B variants during a time-course incubation with Htm1o-Pdi1p. Data shown are the mean values ± one s.d. from experimental triplicates.

Analysis of the demannosylation of RNase B variants by Htm1p-Pdi1p revealed that Htm1p-Pd1ip was able to distinguish differences in conformations of these variants (Fig. 6*C*). Htm1p-Pdi1p showed minimal mannosidase activities on the native-like RNase BS, suggesting that Htm1p-Pdi1p is insensitive to the mild increase in peptide flexibility around the N-glycosylation site of RNase BS [41]. For nonnative variants, we observed a clear trend of increasing demannosylation efficiency as one goes from the globally unfolded Carb-RB, to the more compact but heterogeneous Scr-RB, and finally to the most compact and homogenous form RBsp. Demannosylation of Carb-RB continued during prolonged incubation (Fig. S10*A*), indicating that the different levels of demannosylation at earlier time points resulted from slower kinetics. Scr-RBsp and Carb-RBsp were similarly susceptible to Htm1p-Pdi1p as their full-length counterparts, suggesting that the difference in the kinetics was not due to the absence of the S peptide *per se* (Fig. S10*B*). The slower kinetics against Carb-RB was not due to alkylation of cysteines, as a fully reduced RNase B (Red-RB) was demannosylated at a rate similar to Carb-RB (Fig. S10*C*). In addition, Htm1p-Pdi1p was not inhibited when its cysteines were first blocked with iodoacetamide before reaction with RBsp, ruling out the possibility of a cysteine-mediated demannosylation mechanism (Fig. S10*D*). To conclude, our findings demonstrate that not only does Htm1p-Pdi1p differentiate nonnative proteins from native ones, but it also prefers nonnative proteins with compact, partially structured conformations over globally unfolded, supporting our latter model that Htm1p-Pdi1p preferentially targets nonnative proteins at specific folding states.

## Discussion

A fundamental question in the ERAD field is what are the biochemical properties that differentiate an ERAD substrate from a normal protein in the ER [3, 20]. The unique requirement of a N-glycan remodeling step for ERAD-L commitment potentially provides a chemical handle to investigate the biochemical basis that determines an ERAD-L substrate. Here, our study provides biochemical evidence to support a model that this commitment step by Htm1p-Pdi1p is closely coordinated with the conformations of potential ERAD-L substrates. As such, the resulting terminally exposed α1,6-linked mannose is suitable to be a folding-state mark that flags folding defects in the attached proteins.

Several lines of biochemical evidence support our conclusion that Htmi1p-Pdip is a folding-sensitive mannosidase. First, the mannosidase activity of Hm1p-Pdi1p against misfolded forms of CPY and RNase B variants but not free Man8 suggests that unlike Mns1p, Htm1p-Pdi1p is a glycoprotein-specific mannosidase. Second, we see a marked enhancement of demannosylation activity of Htm1p-Pdi1p when native CPY and native RNase B were artificially converted to misfolded forms. Third, we see significantly higher rate of activity of Htm1p-Pdi1p against partially structured RNase B variants compared to their globally unfolded forms indicating that Htm1p-Pdi1p can distinguish different nonnative states. Our biochemical analysis of the conformational sensitivity of Hmt1p-Pdi1p is supported by *in vivo* studies revealing a higher abundance of steady-state Man7 on ER-retained CPY* than CPY in wild-type yeast; this suggests that generation of Man7 is not a passive, universal event of prolonged ER retention.

The observed preference for partially folded forms can explain why Htm1p-Pdi1p has higher mannosidase activity against Scr-CPY than Carb-CPY. On the other hand, the absence of ER chaperones that are critical to keep CPY* in partially structured states may hamper Htm1p-Pdi1p from exerting its full activity on these substrates in our system [42]. As proteins populated at partially structured states have the propensity to form aggregates and amyloids [43, 44], this conformational preference of Htm1p-Pdi1p may ensure that proteins on the verge of aggregating are promptly committed for degradation. Additionally, this folding-sensitivity of Htm1p-Pdi1p can ensure that nascent, globally unfolded polypeptides are allowed sufficient time for folding.

Interestingly, this conformational preference is reminiscent of that of UDP-glucose glucosyltransferase (UGGT), which reglucosylates partially structured proteins to help their retention in the ER folding cycle [45]. The presence of multiple N-glycan-remodeling enzymes with a similar preference for compact intermediates further argues for the broad biological importance of detection of partially structured proteins in the ER. The essential interaction of Htm1p with Pdi1p is also reminiscent of the predicted structure of UGGT, which contains three thioredoxin-liked domains that may contribute to peptide binding [46]. The reconstitution system that we have established here provides a roadmap for future structural investigation of whether and how the thioredoxin domains of Pdi1p contribute to the recognition of partially structured proteins. Furthermore, our discovery that Htm1p-Pdi1p can remove the α1,2-linked mannose from Branch C on Man9 without the prior action of Mns1p suggests a future direction to apply synthetic Man9-glycoconjugates that were developed for the investigation of UGGT [47] for more detailed analysis on how the N-glycan is used as a potential time indicator by Htm1p-Pdi1p for glycoprotein quality control in the ER.

To conclude, in addition to the previous characterized ERAD-L surveillance step mediated by the Yos9p-Hrd3p complex [6, 7, 16], here we provide evidence for an upstream folding surveillance step that is mediated by Htm1p-Pdi1p. The presence of multiple folding surveillance steps, each with its own unique N-glycan processing or recognition functions, supports a model that ERAD-L follows a kinetic proofreading mechanism to achieve high fidelity in targeting the “right” proteins for degradation [6, 7, 48].

## Materials and Methods

### Yeast strains

All yeast transformations were conducted according to standard procedures. For chromosomal overexpression of *HTM1* (yJW1819 and yJW1820), the chromosomal *HTM1* locus was deleted from BY4741 by a *URA3* selection marker. Double-stranded DNA starting either with the endogenous 5’UTR promoter or a *TDH3* promoter for overexpression, followed by the full coding-region of wild-type or mutant *HTM1*, with a C-terminal 3xFLAG-HDEL tag, 3’-UTR of *HTM1* and finally a KANMX selection marker was then introduced into the *htm1Δ∷URA3* strain. A pGAL1-MNS1-3xFLAG∷KANMX strain was generated by the same procedure for the preparation of 3xFLAG-tagged Mns1p that is controlled by a *GAL1* promoter. See **Table S1** for the list of strains used in this study.

### Plasmids

The plasmid for *GAL1* promoter-driven CPY* expression (pJW1528) was kindly provided by Tom Rapoport (Harvard Medical School). To generate a native CPY expression plasmid (pJW1529), PCR primers carrying the native G255 codon were used to amplified the G255R region from the CPY* expression plasmid, and then assembled into the same vector backbone by the Gibson method. To generate a CPY*0001 expression plasmid (pJW1530), PCR primers carrying asparagine to glutamine mutations of the first three N-glycosylation sites were used to make mutant fragments on CPY* by PCR amplification, which were then assembled into the same 2-μm vector backbone by the Gibson method [49]. See **Table S2** for the list of plasmids used in this study.

### Antibodies

Anti-FLAG M2 mouse monoclonal antibody and the antibody-conjugated agarose beads were purchased from Sigma-Aldrich (St. Louis, MO). Anti-yeast Pdi1p rabbit antisera was a gift from Peter Walter (University of California, San Francisco), which was confirmed by recombinantly expressed Pdi1p. IR fluorescence-conjugated secondary antibodies were purchased from LI-COR (Lincoln, NE).

### Recombinant preparation of Htm1p-Pdi1p and Mns1p

The yJW1820 yeast was grown in three liters of YEPD 30°C to reach the stationary phase. Cells were harvest by centrifuging at 4,000 x *g* for 5 minutes, washed with cold deionized water, and stored at −80°C. All of the following purification steps were carried out at 4°C. Approximately 15 g of cell pellet was mixed with 5 g of dry ice, and then was ground by a Proctor Silex E160BY grinder (Southern Pine, NC) for a 30-second On/Of cycle for 5 times. The cell ground was vacuum-degassed for 5 minutes to remove residual dry ice, and thawed and resuspended in 25 ml Lysis Buffer (20 mM HEPES pH 7.0, 150 mM NaCl, 2 mM CaCl_2_, 0.5% (v/v) NP-40, 15% (v/v) glycerol, 1x *cOmplete* EDTA-free protease inhibitor cocktail (Roche Diagnostics)). The slurry was centrifuged at 6,000 x *g* twice to remove cell debris, nutated for 30 minutes to solubilize the lipid membrane, and then ultracentrifugation at 60,000 x *g* for 30 minutes. The supernatant was collected and transferred into a new tube containing approximately 300 μl bed-volume of anti-FLAG M2 affinity agarose beads (Sigma-Aldrich) that has been equilibrated with the Lysis Buffer. The mixture was nutated for 3 hours and then transferred into an open column. The column was washed with 3 ml Wash Buffer 1 (20 mM HEPES pH 7.0, 150 mM NaCl, 2 mM CaCl_2_, 1x *cOmplete* EDTA-free protease inhibitor cocktail, 15% (v/v) glycerol), 1 ml Wash Buffer 2 (20 mM HEPES pH 7.0, 300 mM NaCl, 2mM CaCl_2_, 15% (v/v) glycerol), and 1 ml Wash Buffer 3 (20 mM HEPES pH 7.0, 150 mM NaCl, 2 mM CaCl_2_, 20 mM imidazole, 15% (v/v) glycerol). To release Htm1p-Pdi1p from the affinity beads, the column was closed by an end-cap, and 10 μg TEV protease and 300 μl Wash Buffer 3 were added. After overnight incubation without agitation, the cleaved products were eluted with 400 μl more Wash Buffer 3. All eluent was collected into a bottom-sealed Micro Bio-Spin column (Bio-Rad, Hercules, CA) containing 100 μl bed volume of Ni-NTA agarose beads (Qiagen, Valencia, CA) that had been equilibrated with Wash Buffer 3. After 30 minutes of incubation, the spin column was centrifuged at 50 x *g* for 10 seconds. The eluent was collected, and the Ni-NTA beads were further eluted with 100 μl Wash Buffer 3 twice more for full elution. The eluent was buffer-exchanged into HSCG Buffer (20 mM HEPES pH 7.0, 150 mM NaCl, 2 mM CaCl_2_, 15% (v/v) glycerol) on a PD MiniTrap G-25 column (GE Healthcare Life Sciences, Pittsburgh, PA). Purified Htm1p-Pdi1p was stored in aliquots at −80°C. Endoglycosidase H (New England Biolabs, Ipswich, MA) treatment was carried out according to the manufacturer’s protocol. For size-exclusion column chromatography, Htm1p-Pdi1p was eluted with 500 μl of 1mg/ml 3xFLAG peptide (Sigma-Aldrich) in Wash Buffer 3, instead of TEV protease. The eluent was then loaded and separated in a Superdex 200 10/300 GL column (GE Healthcare Life Sciences) for subsequent western blot and mannosidase assay. Protein concentration was determined by Pierce BCA protein assay (Thermo Fisher Scientific, Waltham, MA).

For Mns1p preparation, yJW1832 was grown in 3-liter dextrose-free YP with 2% (w/v) raffinose at 30°C to early stationary phase. Galactose was then added to 2% (w/v) to induce expression for 12 hours. Cells were then harvested. Cell lysis and affinity purification were carried with the same procedure as Htm1p-Pdi1p.

### Recombinant preparation of CPY and CPY*

CPY plasmids were introduced into the desired yeast strain, and the transformed cell was grown in 3-liter SC-LEU with 2% (w/v) raffinose till OD_600_ reached 1.0~1.2. Galactose was then added to 2% (w/v) to induce expression at 30°C for another 12 hours. Cells were then harvested and stored at −80°C. All of the purification steps were carried out at 4°C. For CPY*, 15 g of cells were ground and then resuspended in 25 ml C-Lysis Buffer (20 mM HEPES pH 7.0, 500 mM NaCl, 10 mM imidazole, 10 mM TCEP-HCl, 6 M guanidine hydrochloride, 1% (v/v) NP-40, 1x *cOmplete* protease inhibitor cocktail). The slurry was nutated for 30 minutes to solubilize the inclusion body, centrifuged at 6,000 x *g* for 10 minutes, and then ultracentrifuged at 60,000 x *g* for 30 minutes. Supernatant was collected and loaded on a Ni-NTA column containing 1 ml bed-volume of resins that had been equilibrated with C-Lysis Buffer. The column was washed with 10 ml C-Wash Buffer (20 mM HEPES pH 7.0, 500 mM NaCl, 10 mM imidazole, 10 mM TCEP-HCl, 6 M guanidine hydrochloride), and then eluted with C-Elution Buffer (20 mM HEPES pH 7.0, 500 mM NaCl, 500 mM imidazole, 6 M guanidine hydrochloride). The eluent was buffer-exchanged into HSGG Buffer (20 mM HEPES pH 7.0, 150 mM NaCl, 15% (v/v) glycerol, 0.1% (w/v) octyl-α-glucopyranoside) and stored at −80°C in aliquots.

The purification of native CPY was performed in the same way as CPY* except that guanidine hydrochloride and TCEP-HCl were omitted from C-Lysis and C-W Buffer, respectively. To make Scr-CPY, CPY was first reduced with 6 M guanidine hydrochloride and 5 mM dithiothreitol (DTT) at 42°C for one hour. N,N,N’,N’-tetramethylazodicarboxamide (diamide) (Santa Cruz Biotechnology, Dallas, TX) was then added to a final concentration of 25 mM. The solution was kept at room temperature for one hour and then buffer-exchanged into HSGG buffer. To make Carb-CPY, CPY was first reduced with 6 M guanidine hydrochloride and 5 mM DTT at 42°C for one hour. Iodoacetamide was then added to 25 mM. The solution was kept dark at room temperature for one hour and then buffer-exchanged into HSGG buffer. Ellman’s reagent was used to check the status of cysteines.

### Preparation of RNase B

Crude bovine pancreatic RNase B (Sigma-Aldrich) was first enriched for the Man_8_GlcNAc_2_-abundant species. Generally, 50 mg of crude RNase B was dissolved in EQ Buffer (20 mM HEPES pH 7.0, 150 mM NaCl, 5 mM CaCl_2_, 5 mM MgCl_2_, 5 mM MnCl_2_) in 10 mg/ml concentration, and then mixed with 5 ml bed volume of EQ Buffer-equilibrated concanavalin A Sepharose beads (Sigma-Aldrich) at 4°C for 2 hours. The slurry was subsequently transferred into an open column, washed sequentially with 15 ml EQ Buffer, 50 ml High-Salt Wash Buffer (20 mM HEPES pH 7.0, 500 mM NaCl, 1 mM CaCl_2_, 1 mM MgCl_2_, 1 mM MnCl_2_), 10 ml EQ Buffer, and finally 200 ml Low-Glc Wash Buffer (20 mM HEPES pH 7.0, 150 mM NaCl, 50 mM Methyl-α-*D*-glucopyranoside). The column was first eluted 100 ml 200mM-Glc Buffer (20 mM HEPES pH 7.0, 150 mM NaCl, 200 mM Methyl-α-*D*-glucopyranoside) into 20 ml/fraction, and further eluted with 100 ml 1M-Glc Buffer (20 mM HEPES pH 7.0, 150 mM NaCl, 1 M Methyl-α-*D*-glucopyranoside). The N-glycan profile of RNase B from each step were checked by MALDI-TOF MS. Fractions containing the desired glycan species were pooled together, adjusted to pH 4.0 by acetic acid, and then loaded onto a 5-ml HiTrap-SP FF cation-exchange column (GE Healthcare Life Sciences) that had been equilibrated with Acidic Buffer (20 mM HEPES pH 4.0, 150 mM NaCl). The cation-exchange column was washed with 25 ml Acidic Buffer, and then eluted with Neutral Buffer (20 mM HEPES pH7.0, 500 mM NaCl). The eluent was buffer-exchanged into HSG Buffer by a PD10 desalting column (GE Healthcare Life Sciences). Purity was checked by size exclusion column chromatography.

For the preparation of RBsp, 1 mg RNase B was mixed with 20 μg freshly dissolved subtilisin (Sigma-Aldrich) in 2 ml HSG buffer. The mixture was kept at 4°C overnight, and 20 μg more subtilisin was added for another one-hour incubation at 4°C. The pH of the mixture was then adjusted to 2.0 with 1 M hydrochloric acid for one hour on ice to destroy subtilisin. Trichloroacetic acid (TCA) was then added to 10% (w/v), and the solution was warmed up to room temperature to allow RBsp precipitation overnight. The mixture was centrifuged at 15,000 x *g* for 10 minutes, and the supernatant was removed. The pellet was dissolved with 9 M deionized urea, and then TCA-precipitated again to fully remove residual S-peptide. The pellet was dissolved with 9 M urea and buffer-exchanged into HSGG Buffer and stored at −30°C in aliquots. Disulfide-scrambling and carbamidomethylation of RNase B and RBsp were carried out with the same procedures as for Scr-CPY and Carb-CPY.

### Circular dichroism analysis

RNase B variants were buffer-exchanged into 10 mM sodium phosphate, pH 7.0 to a final concentration of 10 μM. Measurement was conducted on a Jasco J-715 spectrometer in a 1-mm cuvette at 30°C. The spectrum was recorded over the range of 190-250 nm at a scanning speed of 20 nm/min with 1.0 nm bandwidth.

### Limited proteolysis of RNase B variants

Trypsin (Sigma-Aldrich) was added to a final concentration of 50 ng/μl into 10 μM RNase B variants at the beginning of the time-course reaction at room temperature. At each time point, equal amount of sample was withdrawn and mixed with 1/10 volume of phenylmethylsulfonyl fluoride (Sigma-Aldrich) to stop the reaction for reducing SDS-PAGE analysis.

### PNGase F sensitivity of RNase B variants

Glycerol-free PNGase F (New England Biolabs) was added to a final concentration of 0.5 unit/μl into 10 μM RNase B variants at the beginning of the time-course reaction at room temperature. At each time point, equal amount of sample was withdrawn and mixed with 4x SDS sample buffer to stop the reaction for reducing SDS-PAGE analysis.

### Glycan profiling by MALDI-TOF MS

The procedure of N-glycan profiling by MALDI-TOF MS was based on published methods [50, 51] with some modifications. N-glycoproteins were separated by SDS-PAGE and stained by Coomassie Blue R250. Individual gel bands were excised and destained twice with 100 μl 50% (v/v) acetonitrile with 10 mM Na_2_CO_3_ at 50°C for 30 minutes with vigorous shaking. The gels were then reduced by 50 mM DTT in 20 mM Na_2_CO_3_ at 50°C for 30 minutes, and then alkylated by adding iodoacetamide to 150 mM and incubated at dark at room temperature for 30 minutes. The gels were washed twice by 50% (v/v) acetonitrile with 10 mM Na_2_CO_3_ at room temperature for 30 minutes, dehydrated by 100% acetonitrile at room temperature for 10 minutes, and evaporated in a centrifugal evaporator for 10 minutes. To each dried gel piece, 5 units of glycerol-free PNGase F (New England Biolabs) in 20 μl of 20 mM Na_2_CO_3_ were added, and the mixture was incubated at 37°C overnight. To extract the released glycans, 100 μl of ddH_2_O was added, and the mixture was sonicated in the water-bath sonicator for 30 minutes. The solution was collected, and extraction was repeated for two more times. To desalt the pooled extract, each sample was loaded into a 20-μl filter tip filled with 10 mg of graphitized carbon (Grace Davidson Discovery Sciences) that had been sequentially washed with 1 ml acetonitrile and 1 ml of ddH_2_O. After loading the sample, the tip-column was washed with 1 ml of ddH_2_O, and eluted with 100 μl of 25% (v/v) acetonitrile. For free glycans, the reaction solution was directly loaded onto the graphitized carbon tip-column and processed with the same procedure. The eluent was evaporated in the centrifugal evaporator at 60°C. The dried sample was resuspended with 5 μl of ddH_2_O, and 1 μl of the sample was spotted on the MALDI target plate and vacuum dried. The plate was then washed with pure acetonitrile. 1 μl of 5 mg/ml 2.5-dihydroxybenzoic acid (DHB) (Sigma-Aldrich) dissolved in 50%_(v/v)_ acetonitrile was then spotted onto each sample spot and dried. The mass spectrometric analysis was conducted on a Voyager Elite DE-STR Pro mass spectrometer (Applied Biosystems) or a Shimadzu AXIMA Performance mass spectrometer (Shimadzu) in the positive reflectron mode. For data collected in the Voyager mass spectrometer, the spectrum analysis was carried out with the bundled Data Explorer software to extract the values of peak area of the peak matched to each glycan. For spectra collected from the AXIMA Performance mass spectrometer, raw spectra were exported in the mzXML format, and then analyzed by the mMass software [52] to extract the values of peak intensity of the peak matched to each glycan.

### Mannosidase assay

Unless otherwise specified, glycoprotein substrates in defined concentration (2 μM for CPY variants, 10 μM for RNase B isoforms) were mixed with 0.1 μM Htm1p-Pdi1p and/or 0.1 μM Mns1p in Reaction Buffer (20 mM HEPES pH 7.0, 150 mM NaCl, 2 mM CaCl_2_, 0.1% (v/v) NP-40). At each desired time-point, EDTA was added to 25 mM to stop demannosylation reaction. The mixture was separated by reducing SDS-PAGE and then subjected to N-glycan-profiling by MALDI-TOF MS.

### NMR analysis

For the Man8-abundant RNase B control, 2 mg of RNase B was reduced by 10 mM DTT at 95°C for 10min and then let cooled down. For the Htm1p-Pdi1p-treated RBsp, 2 mg of RBsp was first incubated with 50 μg of Htm1p-Pdi1p overnight. To both samples, 1000 units of PNGase F were then added for glycan release at 37°C overnight. The next day, the reaction mixture was loaded onto a 200 mg of graphitized carbon column, washed with 5 ml water, and then eluted with 1.5 ml 25% acetonitrile. The eluent was lyophilized, and then dissolved by D_2_O for lyophilization for two more times. The dried sample was then dissolved by D2O for NMR measure. The ^1^H-NMR was carried out on a 400 MHz Bruker AvanceIII HD 2-channel instrument with a TopSpin v3.5 interface. Spectra were pre-collected using default proton method, calibrated to the reference chemical shift of water (4.79 ppm), and then collected with the presaturation method as described by the manufacturer to suppress the signal from water.

## Author Contributions

Conception and design: Y.-C. L., D.G.F., J.S.W.; Acquisition of data: Y.-C.L.; Analysis and interpretation of data: Y.-C.L; Drafting or revising the article: Y.-C.L., D.G.F., J.S.W.; Supervision: D.G.F., J.S.W.

## Acknowledgements

We would like to thank Tom Rapoport and Alexander Stein (Harvard Medical School) for providing the recombinant CPY plasmids and the detailed purification protocol; Peter Walter for providing anti-Pdi1p antibody; current and former members of the Fujimori lab and the Weissman lab for discussion and laboratory methods; Erin Quan Toyama and Jay Read for the initial tests on Htm1; Shoshana Bar-Nun, Elizabeth Costa, Joshua Dunn, Christina Fitzsimmons, Calvin Jan, Kamena Kostova, Melanie Smith, Lindsey Pack, Dan Santos, and Erin Quan Toyama for critical comments on the manuscript. We thank Yao-ming Huang for the assistance with circular dichroism analysis (Kortemme group, UCSF). We also thank the UCSF Mass Spectrometry Facility, the DeGrado group, and the UCSF NMR laboratory for access to the mass spectrometers and the NMR instrument. We are grateful for the support from the Howard Hughes Medical Institute International Student Research Fellowship (Y.-C.L.), Sandler Foundation and the UCSF Program for Breakthrough Biomedical Research Award (D.G.F. and J.S.W.), the Howard Hughes Medical Institute (J.S.W.), and National Institute of Health (U01 GM098254 to J.S.W.).

## Abbreviations List

ERAD: Endoplasmic reticulum-associated protein degradation
Hex10: Hex_10_GlcNAc_2_
Man7~9: Man_7~9_GlcNAc_2_
CPY: Yeast pro-carboxypeptidase Y
CPY*: CPY G255R mutant
MALDI-TOF MS: Matrix-assisted laser desorption/ionization time-of-flight mass spectrometry
DTT: Dithiothreitol
Diamide: N,N,N’,N’-tetramethylazodicarboxamide
IAA: Iodoacetamide
Scr-CPY: Disulfide-scrambled CPY
Carb-CPY: Cysteine-carbamidomethylated CPY
DMJ: 1-deoxymannojirimycin
E222Q: Htm1 mutant with putative active site residue Glu_222_ mutated to Gln
Δ572-657: Htm1 mutant with truncation of the putative Pdi1p-interacting domain
CPY*1110: CPY* mutant with an N479Q mutation at the fourth N-glycosylation site
CPY*0001: CPY* mutant with the first three N-glycosylation sites mutagenized (N124Q, N198Q and N279Q)
CPYm9: Man9-carrying CPY purified from mns1Δhtm1Δ
Scr-CPYm9: Disulfide-scrambled CPYm9
RNase B: bovine pancreatic ribonuclease B
RBsp: RNase BS protein
RNase BS: Non-covalent complex between the S peptide and RBsp
Scr-RB: Disulfide-scrambled RNase B
Carb-RB: Cysteine-carbamidomethylated RNase B
Red-RB: Fully reduced RNase B
Scr-RBsp: Disulfide-scrambled RBsp
Carb-RBsp: Cysteine-carbamidomethylated RBsp

